# Learning high-dimensional reaction coordinates of fast-folding proteins using State Predictive information bottleneck and Bias Exchange Metadynamics

**DOI:** 10.1101/2023.07.24.550401

**Authors:** Nancy D. Pomarici, Shams Mehdi, Patrick K. Quoika, Suemin Lee, Johannes R. Loeffler, Klaus R. Liedl, Pratyush Tiwary, Monica L. Fernández-Quintero

## Abstract

Biological events occurring on long timescales, such as protein folding, remain hard to capture with conventional molecular dynamics (MD) simulation. To overcome these limitations, enhanced sampling techniques can be used to sample regions of the free energy landscape separated by high energy barriers, thereby allowing to observe these rare events. However, many of these techniques require a priori knowledge of the appropriate reaction coordinates (RCs) that describe the process of interest. In recent years, Artificial Intelligence (AI) models have emerged as promising approaches to accelerate rare event sampling. However, integration of these AI methods with MD for automated learning of improved RCs is not trivial, particularly when working with undersampled trajectories and highly complex systems. In this study, we employed the State Predictive Information Bottleneck (SPIB) neural network, coupled with bias exchange metadynamics simulations (BE-metaD), to investigate the unfolding process of two proteins, chignolin and villin. By utilizing the high-dimensional RCs learned from SPIB even with poor training data, BE-metaD simulations dramatically accelerate the sampling of the unfolding process for both proteins. In addition, we compare different RCs and find that the careful selection of RCs is crucial to substantially speed up the sampling of rare events. Thus, this approach, leveraging the power of AI and enhanced sampling techniques, holds great promise for advancing our understanding of complex biological processes occurring on long timescales.

TABLE OF CONTENT GRAPHIC

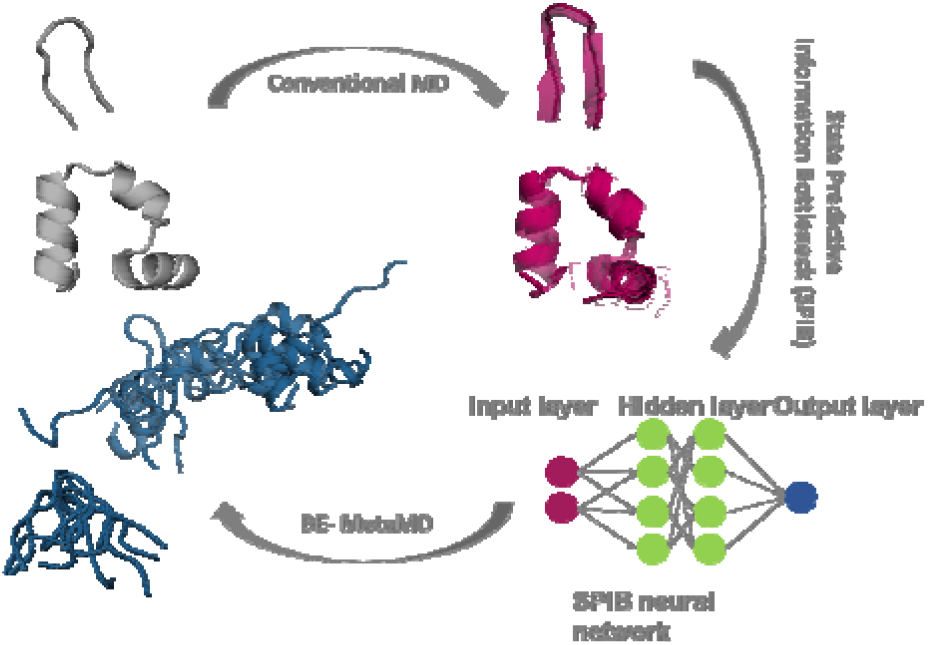

## INTRODUCTION

Protein function is governed by their dynamic nature and is characterized by their ability to adapt different conformations in solution.^1, 2^ Various Nuclear magnetic resonance (NMR), cryo-EM and crystal structures are available up to now, showing that proteins can adopt multiple conformations, undergoing reorientations or transitions from disordered to ordered states.^2, 3^ However, the experimental resolution sometimes does not allow to observe conformational changes at a molecular level to fully understand their mechanism.^4^ As an alternative, MD simulations represent an attractive tool to capture the details of protein conformational dynamics and determine the probabilities of these conformational rearrangements. Many biological processes require transitions between conformational states separated by high energy barriers, which naturally happen at milliseconds timescales or even longer.^5–8^ Unfortunately, overcoming these high energy barriers with conventional MD (cMD) simulations at low computational cost still represents a challenge.^9^ To address this problem, many enhanced sampling algorithms have been developed, such as umbrella sampling^10^, hyperdynamics^11, 12^, metadynamics^13–15^, replica exchange^16^ and many others.^17, 18^ Many of these methods require the selection of appropriate RCs that can comprehensively describe the process of interest.^19^ However, identifying suitable RCs in advance is often a challenging task for numerous systems and complex processes. Although this field substantially advanced in the last decades, the sampling of rare events and the selection of suitable RCs remains challenging.

In recent years, AI has made significant contributions and revolutionized the field of computational chemistry and protein structure prediction. Machine learning (ML) algorithms and deep neural networks revealed their applicability in MD-related fields, enabling efficient modeling of complex biological systems and providing valuable insights into protein folding.^20–22^ Among these methods, the reweighted autoencoded variational Bayes for enhanced sampling (RAVE) involves iterations between molecular simulations and deep learning to obtain physically interpretable RCs and to accelerate the generation of accurate probability distributions, surpassing the capabilities of unbiased MD.^23^ A direct descendant of the RAVE method is the State Predictive Information Bottleneck (SPIB) approach.^24^ This framework aims to find RCs that carry the minimal information about the past but can predict the future state of the system. The physically interpretable RCs learned by SPIB can be used as biasing coordinates in enhanced sampling methods, to identify metastable states and accelerate the conformational sampling. This approach has previously been applied in combination with well-tempered metadynamics^14^, to find the RCs from undersampled trajectories and accelerate the molecular dynamics of two model systems.^25^ While many attempts focused on reducing the RC dimensionality, many practical systems demand high-dimensional RCs for an accurate description of their complex behavior.^26^

In this work, we employ SPIB with a variational autoencoder (VAE)^27^ architecture for learning high-dimensional RCs, implemented through an appropriate enhanced sampling method, namely bias exchange metadynamics simulations (BE-metaD).^28^ In this technique, multiple replicas of metadynamics simulations are performed, each biasing a different set of RCs. The exchanges between the replicas allow an efficient multidimensional sampling of the complex free energy landscape. BE-metaD has already been successfully applied to study the conduction and selectivity in ion channels, to investigate protein folding and to capture conformational rearrangements in antibodies.^28–32^ Although, in principle BE-metaD can implement many RCs, the sampling can be significantly improved through the construction of an optimal RC. This makes the combination of SPIB with BE-metaD very promising as we explore in this following. Here, we apply our protocol to investigate the unfolding mechanism of two small proteins, namely chignolin and villin (**Figure 1**). These are two fast-folding proteins, with mean folding times of respectively 0.6 and 2.8 µs and mean unfolding times of respectively 2.2 and 0.9 µs.^33^

**Figure 1.**
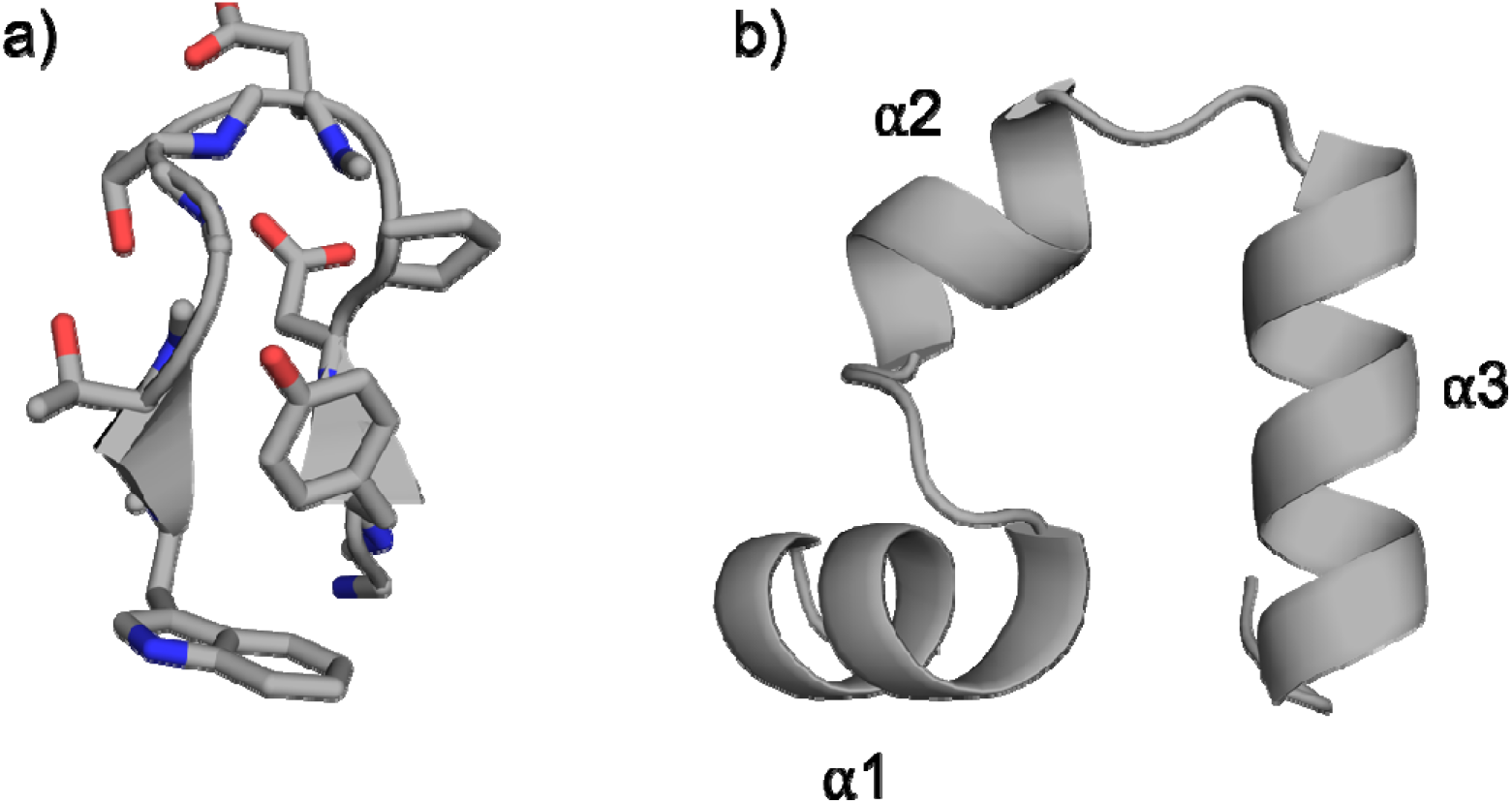
Illustration of the analyzed systems: a) chignolin, a 10-residues peptide; b) villin, a peptide composed of 35 residues.

Chignolin is an artificial peptide composed of 10 residues (GYDPETGTWG) that folds into a β-hairpin structure in water (**Figure 1a**).^34, 35^ The villin headpiece is instead a 35 residues peptide, that folds into three alpha helices (α1, α2 and α3) with a hydrophobic core composed of three phenylalanines (**Figure 1b**).^36, 37^

For these two systems, we applied SPIB to learn the RCs to enhance the conformational sampling from short cMD simulations. The resulting RCs were then used as collective variable for the BE-metaD simulations. The integration of SPIB with BE-metaD simulations allowed to identify suitable RCs, that significantly accelerated the sampling of the conformational transitions.

## RESULTS

We ran relatively short unbiased simulations (200-1000 ns) of two proteins - chignolin and villin. Subsequently, we applied the SPIB algorithm for both systems to train and identify appropriate RCs capable of capturing the conformational transitions in solution involved in the unfolding process. SPIB was trained separately on the ψ backbone torsion angles and on the pairwise distances between the centers of mass (COMs) of the heavy atoms of each residue. The 6 RCs output of SPIB of each trained model were then used as collective variables for the 6 replicas of BE-metaD simulations. In the following, we compare the resulting ensemble from different input features and how SPIB in combination with BE-metaD can accelerate the sampling of the proteins’ unfolding process.

### Unfolding of Chignolin

For chignolin, we followed two independent approaches to train SPIB. Firstly, on the ψ backbone torsion angles and secondly, on the pairwise distances between the COMs of the heavy atoms of each residue. To this end, we calculated these descriptors for a 1 µs and for a 200 ns of cMD simulation. In both, the 200 ns and 1 µs cMD simulation, chignolin is mostly trapped in the folded state. Using the resulting 6 RCs obtained from SPIB, we ran BE-metaD simulations. The result using a linear encoder in the feed-forward SPIB VAE architecture starting from the 1 µs cMD simulation is shown in **Figure 2**. We use a continuous 6 µs cMD simulation as reference simulation, which extensively samples the unfolding process of chignolin and we compare it, using a PCA analysis, to the space sampled in the biased simulations. This representation allows a direct comparison of the free energy landscapes sampled by the unbiased and the biased simulations.^38, 39^ All the reweighted PCA plots shown in **Figure 2** have been transformed to the same coordinate space. **Figure 2c** shows that 100 ns of BE-metaD simulations can cover a similar space compared to the long cMD and thoroughly sample the unfolding process of chignolin. This is true for the RCs learned from the distance features and for the torsion features.

**Figure 2.**
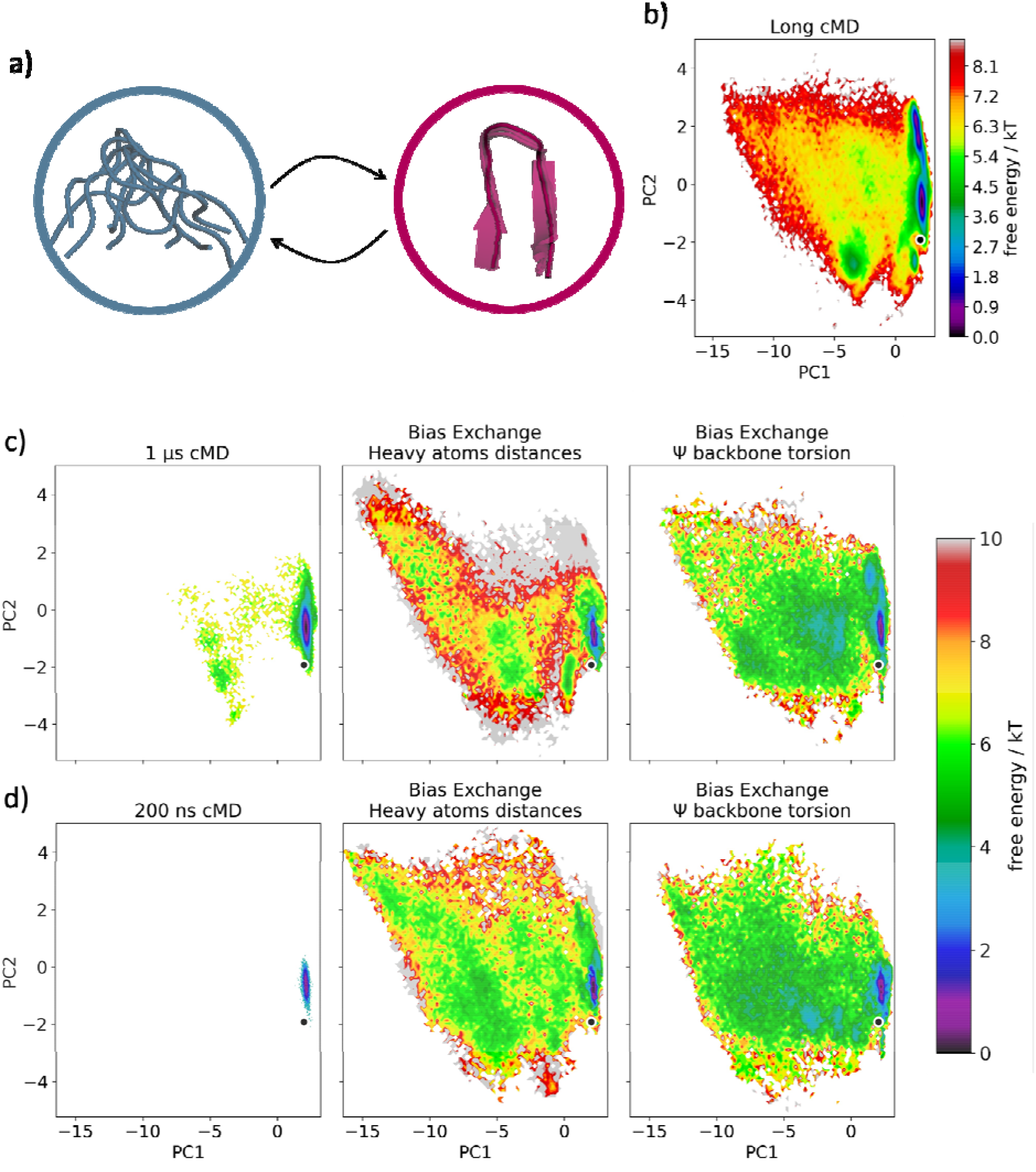
Unfolding process of chignolin. **a)** Structural representations of the two conformational ensembles during chignolin unfolding: on the right, the folded state, on the left, the unfolded one. **b)** PCA space obtained from the 6 µs cMD simulation**, c)** and **d)** Comparison of the PCA spaces (in the same coordinate space) obtained from the short cMD simulations from where the features for SPIB training were calculated, and the 100 ns BE-metaD. The central plots in **c)** and **d)** show the resulting free energy space after training SPIB on the distances between COMs of heavy atoms calculated in the first 1 µs and 200 ns of cMD, respectively. For the plots on the right side, instead, SPIB was trained on the Ψ backbone torsions calculated in the first 1 µs and 200 ns of cMD, respectively. In all the cases, SPIB was trained using a linear encoder.

In addition, we trained SPIB on the features calculated from the 200 ns of cMD trajectory. Also in this case, the unfolding process can be captured in only 100 ns BE-metaD simulations, resulting in similar findings compared to the BE-metaD simulations based on the 1 µs cMD training (**Figure 2d**). Again, the difference between the distance and torsion features used to train the SPIB model is negligible. The results of the trajectories using a non linear encoder with ReLU activation function are shown in **Figure S1**. In this case, differences in the sampled conformational space can be observed depending on the features used for the SPIB training. In fact, the BE-metaD simulations based on RCs derived from training on the 1 µs cMD simulation distance features exhibit a broader conformational space compared to those trained on backbone torsions (**Figure S1a**). The performance decreases when SPIB is trained using the non linear encoder on the features obtained from only 200 ns of cMD (**Figure S1b**).

### Unfolding of villin

As described above for chignolin, also for villin we trained SPIB separately on the distance and angle features calculated for the first microsecond of the available continuous 10 µs cMD simulation. No partial unfolding event is captured in the 10 µs cMD simulation. However, the 100 ns BE-metaD simulations reveal a much larger conformational space including the partial unfolding of the peptide. **Figure 3** shows a comparison of the reweighted PCA spaces in the same coordinate system using non linear and linear encoders, including the 10 µs cMD as reference. In the left plots in **Figure 3d** and **3e**, linear encoders are used, whereas the right ones show the results after training a non linear model. In general, the RCs obtained from training SPIB on the distances between the COMs of the heavy atoms of each residue capture the unfolding of villin more thoroughly. This is evident from the BE-metaD simulations, where these RCs as collective variables result in a larger sampling of conformational space. In addition, the models based on linear encoders performed better than the non linear ones, not only in terms of sampling efficiency but also in terms of computational time and costs.

**Figure 3.**
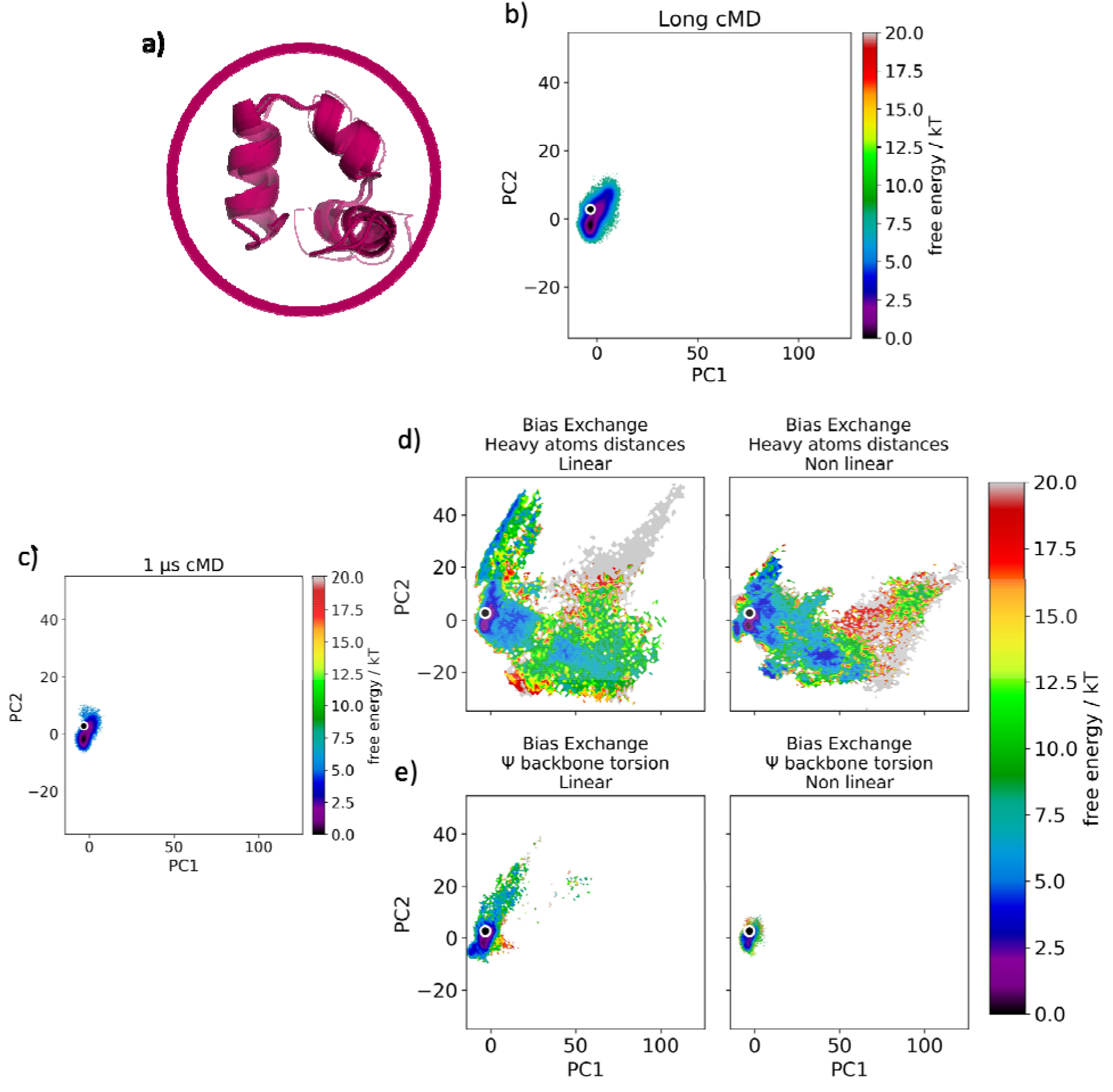
Unfolding process of villin. **a)** Structural representations of a conformational ensemble captured from the 10 µs cMD simulation **b)** PCA space obtained from the 10 µs cMD simulation**, c)** PCA space obtained from the first 1 µs of cMD simulation, used to calculate the features to train SPIB**, d)** and **e)** Comparison of the PCA spaces (in the same coordinate system) obtained from the 100 ns BE-metaD. In **d)** SPIB was trained on the distances between COMs of heavy atoms calculated in the first 1 µs of cMD, using a linear and non linear encoder, respectively. In **e)**, instead, SPIB has been trained on the Ψ backbone torsions calculated in the first 1 µs of cMD, using a linear and non linear encoder, respectively.

### Structural changes involve in the unfolding process

In order to monitor and quantify structural changes of the proteins during the unfolding process, we calculated the secondary structure, the RMSD and the native contacts for the BE-metaD simulations. The six replicas were reordered according to the accepted exchanges and concatenated. **Figure 4** shows the time evolution of the secondary structure elements, the similarity to the native conformation (C𝛼-RMSD) and the fraction of native contacts for the chignolin and villin through the concatenated replicas. The RCs for the considered simulations are derived from training SPIB on a combination of the distances between the heavy atoms COMs of the 1 µs cMD simulation, using a linear encoder. The results for the remaining BE-metaD simulations are shown in **Figure S2**. The RMSD and the native contacts analyses suggest the presence of folding and unfolding events during the simulations (**Figure 4b** and **4d**). The alpha helices α1 and α2 of villin show major secondary structure fluctuations, whereas α3 appears to be the most stable one (**Figure 4c**).

**Figure 4.**
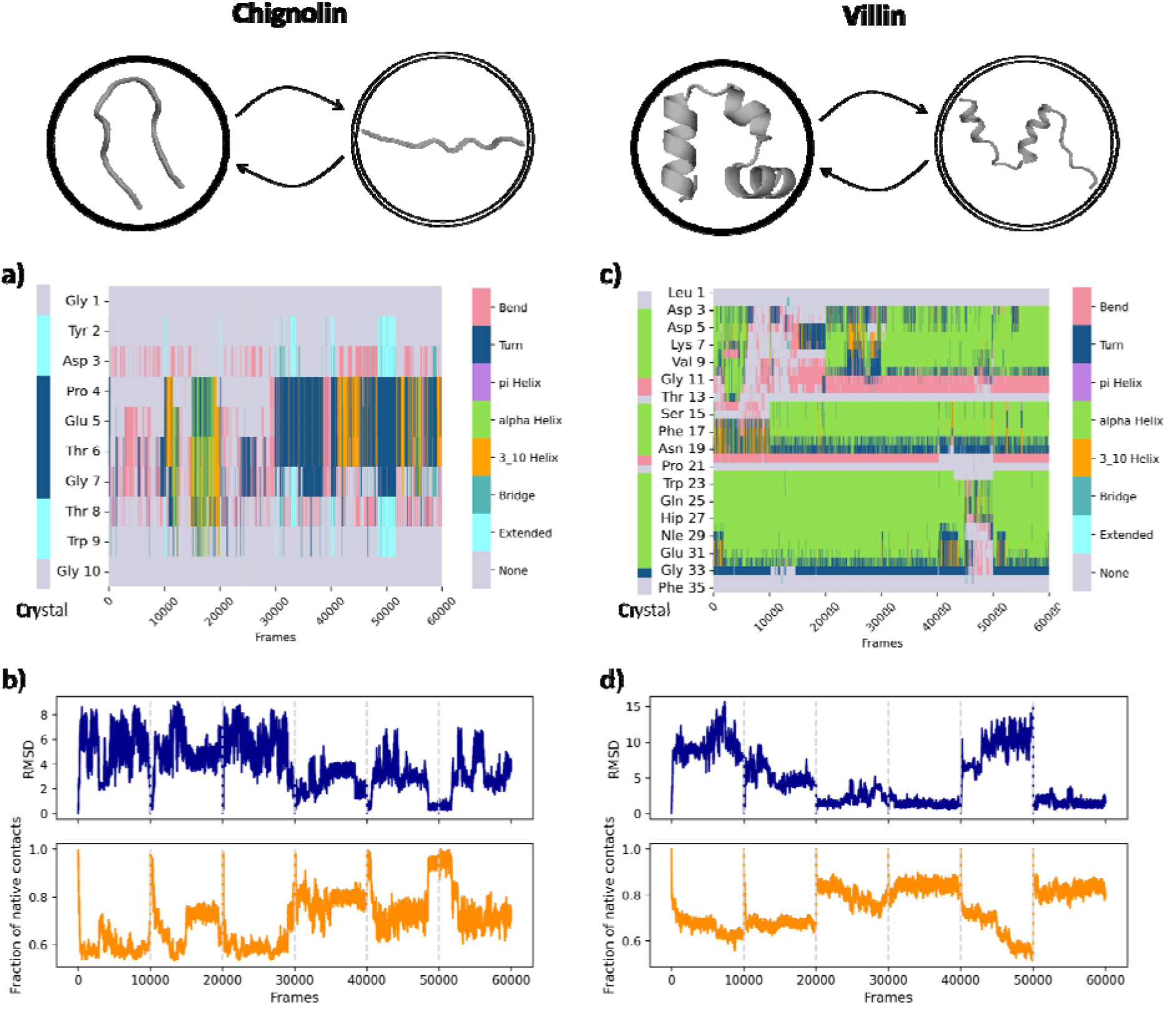
Simulation analyses. Secondary structure (**a)** and **c)**), RMSD and native contacts analyses (**b)** and **d)**) of the BE-metaD simulations of chignolin (left side) and villin (right side). As a reference, the secondary structure of the crystal of the proteins is shown (**a)** and **c)**). The RCs were obtained from SPIB with a linear encoder, trained on the distances between heavy atoms COMs calculated on the first microsecond of the cMD simulations.

## DISCUSSION

In this work, we apply a previously developed ML algorithm – SPIB^24^ – to learn RCs that enhance conformational sampling, and efficiently accelerate the unfolding process of two proteins: chignolin and villin. Due to their small size (10 and 35 residues) and their different folding states in solution, these proteins represent ideal model systems to evaluate the performance of novel and advanced sampling techniques.^39–41^ In addition, they are considered ultrafast folders, since they can adapt the folded conformation in the microsecond timescale, without additional formation of disulfide bridges.^42^ Sampling of the unfolded state of proteins is as crucial as characterizing the folded one, to enable a comprehensive understanding of equilibrium and to provide insights into the protein folding mechanism.^39, 43^ SPIB provides invaluable advantages in combination with enhanced sampling techniques to advance the understanding of complex systems. Even though cMD simulations allow the description of conformational changes that determine the protein function at an atomistic level, they are sometimes limited in capturing the long timescale movements of biomolecular processes.^4, 44^ Hence, many enhanced sampling techniques have been established to overcome this limitation and accelerate the sampling of the conformational space. Especially replica exchange molecular dynamics simulations have proven their applicability for peptides and protein simulations, resulting in many folding events that agree with experimental observations.^45, 46^ A drawback of these techniques is the requirement of RCs that can properly describe the process of interest. Finding such RCs remains often challenging.

SPIB improved the sampling process in two main aspects: a) it simplified the choice of RCs, by finding the most suitable combination from numerous initial features; b) the synergy with BE-metaD simulations drastically accelerated the sampling of the unfolding processes. In fact, the results show that only 100 ns of BE-metaD simulations are sufficient to sample a large conformational space in which transitions between the folded and unfolded states are observed. As previously demonstrated with metadynamics simulations^25^, here we show that also the combination of SPIB with BE-metaD simulations facilitates the choice of RCs and results in a general improvement in performance. In principle, this approach could be transferable also to other methods that can incorporate multiple collective variables, e.g. parallel bias metadynamics.^47^

SPIB was trained on data derived from undersampled cMD trajectories for each system, describing the residue-residue heavy atom COMs distances and the ψ backbone torsion angles for each residue. The resulting RCs were then chosen as collective variables for the BE-metaD simulations. The performance of the method was evaluated with a PCA analysis that compares the sampled space of the BE-metaD simulations with the unbiased simulations. For both proteins, the combination of SPIB with BE-metaD simulations allows a proper sampling of the unfolding process, showing unfolding events. The free energy landscapes of each peptide show a deep energy well corresponding to the native state. Generally, the unfolding can be approximated as a two-state process, in line with the one-dimensional free energy profiles observed in previous folding simulations.^33^

We also compared the variations in performance depending on the initial feature selection. While chignolin shows no noticeable differences considering distance or torsion features, it appears that the unfolding process of villin is more accurately captured by RCs derived from distance features. In contrast, biasing a combination of torsion angles results in a narrower conformational space. This is in line with previous findings for villin that showed that, backbone torsions account accurately for the native state, while contacts in the tertiary structure are an appropriate descriptor to characterize the unfolded region of villin.^41^

Furthermore, we compared the performances of the SPIB models implemented through a VAE architecture with a linear and a non linear encoders respectively. It is known in general, that non linear encoders are highly expressive and therefore, it is recommendable for highly complex systems or processes.^25^ Interestingly, when working within a data-sparse regime, this high expressivity can become problematic due to the model learning noise by overfitting model parameters. This was observed in the case of villin, where the SPIB model with non linear encoder performed poorly compared to the linear encoder model. On the other hand, we observed better sampling performance with a linear encoder, which is also advantageous for the shorter calculation time. In addition, since the SPIB models were trained using normalized data (min-max scaling), for linear encoders, the model weights associated with individual features can be directly used to interpret our RCs and evaluate the contribution of the initial features to the final RCs. It should be noted that in recent literature, interpretation methods e.g, TERP^48^, has been proposed for explaining predictions from black-box AI used in MD simulations (e.g, SPIB models with non linear encoder explored in this research work). By integrating such an interpretation protocol, we can exploit the highly expressive nature of advanced AI models and at the same time ensure that the model is learning system behavior correctly.

In order to prove the validity of the sampled conformations, we performed secondary structure, RMSD and native contacts analyses. Our simulations are in good agreement with previous findings. In fact, the terminal helices of villin, and especially the α3-helix, are more native-like than the middle one.^39, 46, 49^ Chignolin, instead, undergoes variations in the secondary structure during unfolding, mainly forming a bend in the secondary structure and rarely an α-helix or 3_10_- helix, before completely unfolding to an extended structure.^50^

In conclusion, we established a protocol that allows to accelerate the sampling of rare events, such as the protein folding process. While enhanced sampling techniques are already effective in studying intricate biological processes, they typically demand a comprehensive understanding of the system. Here, we propose a protocol that facilitates the choice of suitable RCs, only starting from a short cMD simulation, and use them as collective variables in a BE-metaD simulation. The conformational space and the respective conformations resulting from combining SPIB with BE-metaD describe accurately the unfolding processes of the studied proteins and show good agreement with previous simulations of villin and chignolin in literature.^39, 46, 49, 50^ This work contributes to overcoming the timescale limitation in cMD simulations and thereby advance the understanding of complex systems.

## METHODS

### Structure Preparation and Molecular Dynamics Simulations of Proteins

Experimental structures were available for both our proteins (PDB codes: 1UAO and 2F4K^51^, for chignolin and villin respectively). The AMBER ff14SB force field was used to describe the proteins.^52^ The villin simulation setup followed a previously published protocol.^39^ Two norleucine residues were present in the studied variant of villin, whose parameters were derived using the antechamber module from AmberTools 19.^53^ These residues were capped to avoid charge artifacts. The tLEaP program of AmberTools 19^53^ was used to soak the proteins in cuboidal boxes with a 10 Å protein-wall distance, using TIP3P water model^54^. After an elaborate equilibration protocol^55^, the systems were simulated with the graphical processing unit (GPU) implementation of the pmemd module of AMBER 18^53^, with a timestep of 2 fs. The Langevin thermostat was applied to maintain a constant temperature of 300 K for villin and 300 K for chignolin.^56^ Constant atmospheric pressure was maintained with the Berendsen barostat.^57^ The long range electrostatic interactions were treated with the particle mesh Ewald method.^58^ All bonds including hydrogen atoms were restrained with the SHAKE algorithm.^59^ Villin simulation was run for a total of 10 µs, whereas the chignolin one was run for a total of 6 µs, collecting frames every 10 ps.

### State Predictive Information Bottleneck and Bias Exchange Metadynamics Simulations

SPIB architecture is constructed from an encoder-decoder pair. In this paper we tested performance of linear and non linear encoder respectively. The decoder, instead, is always non linear. SPIB algorithm was trained based on features calculated on only the first microsecond, or the first 200ns of each trajectory, respectively. The ψ dihedral angle of each residue (in rad) and the pairwise distance between the heavy atoms COM of each residue were calculated. The pairwise distances between neighboring residues were not included. SPIB was trained separately on the angles and the distances features. The total number of features is described in **Table S1**. The distances calculation in the villin system resulted in 561 total features. In order to reduce the number of features to obtain the final RCs, a first round of SPIB using a linear encoder was used as a feature selection mechanism to identify the 20 most relevant distances. Consequently, a second round of SPIB was run to obtain the final RCs.

The timeseries of the features obtained from the relatively short unbiased simulations were used to train the encoder – linear or non linear- to convert the high dimensional data into 6 RCs, each used as a biasing variable in one of the replicas of the BE-metaD simulations. The parameters settings chosen for the SPIB trainings are shown in **Table S2**. The simulations were performed at 300 K, in an NpT ensemble with the software GROMACS (version 2020.2)^60, 61^, using plumed implementation (version 2.6.1)^62^. We used a Gaussian height of 1.5 kcal/mol and a bias factor of 10. The Gaussian deposition occurred every 1000 steps. 100 ns of BE-metaD simulations with six replicas were performed for each system. Afterwards, the six replicas were reordered according to the accepted exchanges leading to structurally continuous trajectories. A PCA analysis was used to compare the space sampled by the cMDs and the one obtained from the BE-metaD trajectory. It was performed using the python library PyEMMA 2, with the backbone atoms distances as features.^63^ The PCA space was reweighted following a previously established protocol.^64^

The RMSD, secondary structure and native contacts of the BE-metaD trajectories were calculated by using cpptraj and pytraj from AmberTools 19.^53^ The native contacts are determined by a simple distance cutoff: we used the default cutoff value of 7 Å and the crystal structures as reference. Therefore, any atoms which are closer than the cutoff in the crystal structures are considered to form a native contact.

## ACKNOWLEDGMENTS

N. D. P. has received funding from the European Union’s Horizon 2020 research and innovation programme under the Marie Skłodowska-Curie grant agreement number 847476. The views and opinions expressed herein do not necessarily reflect those of the European Commission. P.T. was an Alfred P. Sloan Foundation fellow during preparation of this manuscript. This work was supported by Austrian Science Fund P34518 and the Austrian Academy of sciences APART-MINT postdoctoral fellowship to M.L.F.Q. We acknowledge CHRONOS for awarding us to access to Piz Daint at CSCS, Switzerland and EuroHPC Joint Undertaking for awarding us access to Karolina at IT4Innovations, Czech Republic.

## DATA AND SOFTWARE AVAILABILITY STATEMENT

The structures for chignolin and villin used in this manuscript are publicly available, with the PDB codes: 1UAO, 2F4K, respectively. The input files to set up the bias exchange metadynamics simulations are available on Github (https://github.com/monilisa2010/SPIB_BE-MetaD.git).

## ASSOCIATED CONTENT

Supporting Information.

## AUTHOR CONTRIBUTIONS

The manuscript was written through contributions of all authors. All authors have given approval to the final version of the manuscript.

## ABBREVIATIONS

MD: molecular dynamics
RC: reaction coordinate
AI: artificial intelligence
SPIB: State Predictive Information Bottleneck
BE-metaD: Bias Exchange Metadynamics
cMD: conventional MD
ML: machine learning
RAVE: reweighted autoencoded variational Bayes for enhanced sampling
VAE: variational autoencoder
COM: center of mass.

## Notes

### Competing Interest Statement

The authors have declared no competing interest.

