## Supporting Information for "Learning high-dimensional reaction coordinates of fast-folding proteins using State Predictive information bottleneck and Bias Exchange Metadynamics"

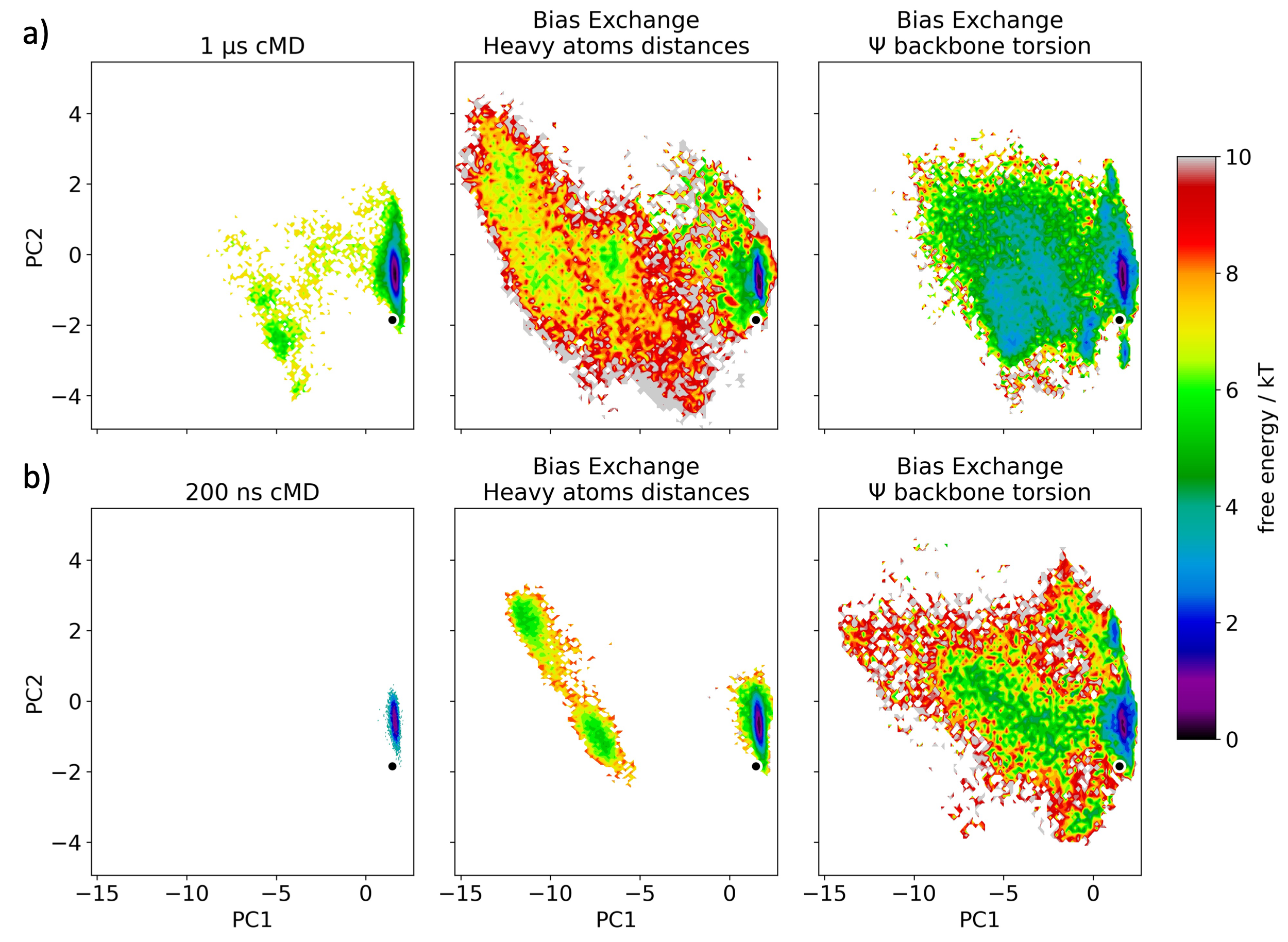

**Figure S1. Unfolding process of chignolin – SPIB performance with non linear encoder a)** and **b)** Comparison of the PCA spaces (in the same coordinate space) obtained from the the short cMD simulations from where the features for SPIB training were calculated, and the 100 ns BE-metaD. The central plots in **a)** and **b)** show the resulting free energy space after training SPIB on the distances between COMs of heavy atoms calculated in the first 1 µs and 200 ns of cMD, respectively. For the plots on the right side, instead, SPIB was trained on the Ψ backbone torsions calculated in the first 1 µs and 200 ns of cMD, respectively. In all the cases, SPIB was trained using a non linear encoder.

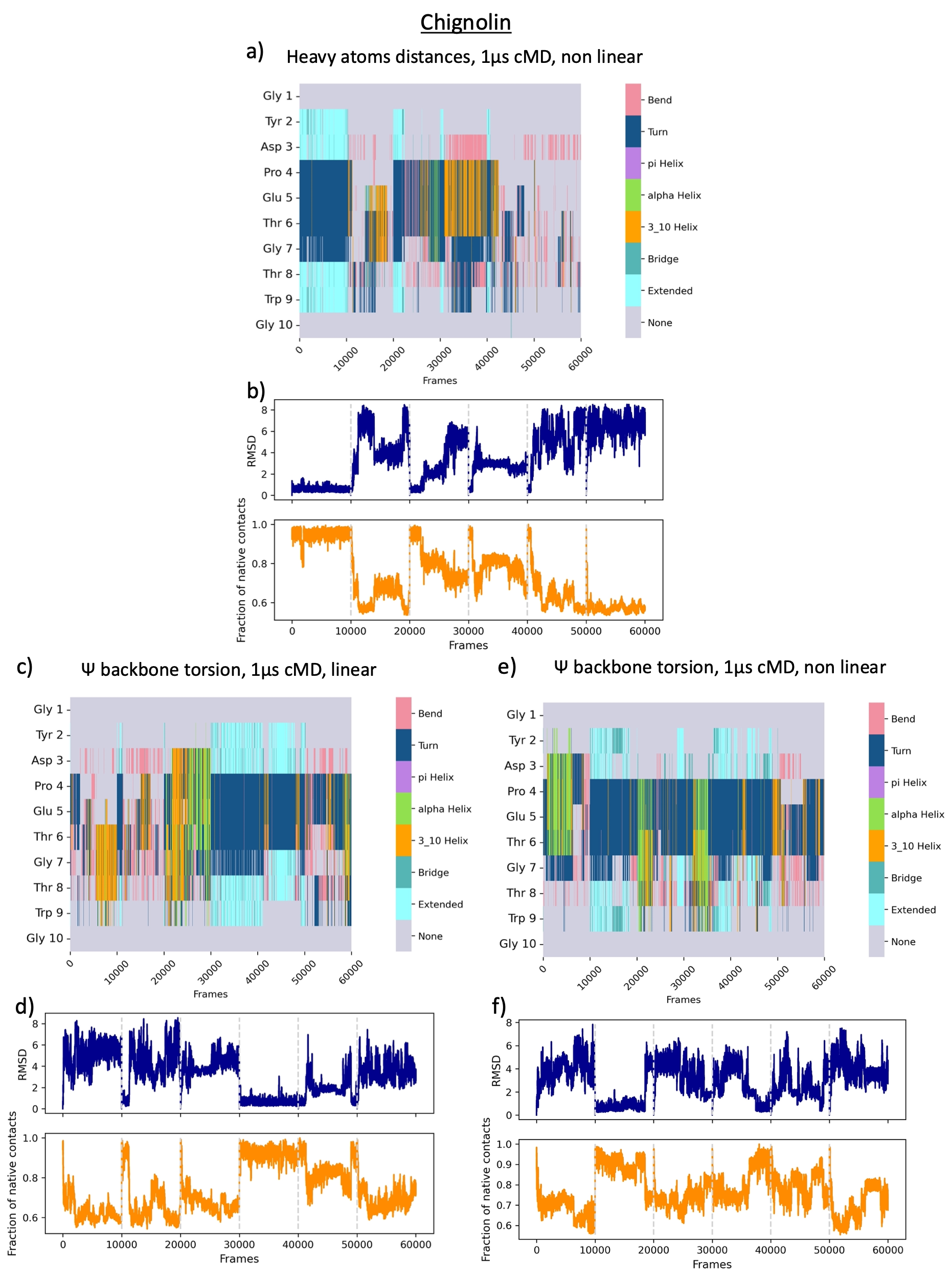

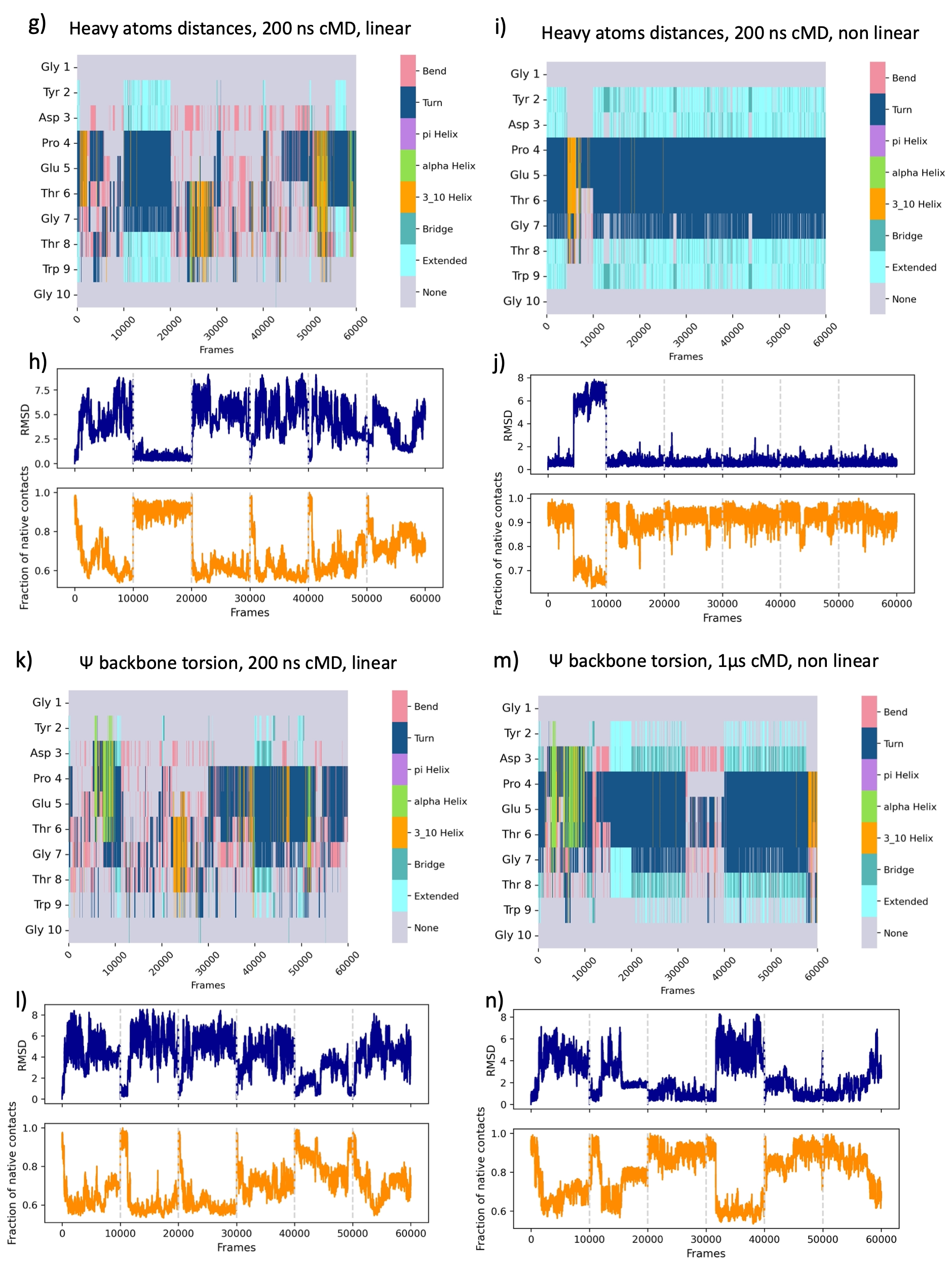

**Figure S2. Analyses of chignolin simulations.** Secondary structure (**a)**, **c)**, **e)**, **g)**, **i)**, **k)** and **m)**), RMSD and native contacts analyses (**b)**, **d)**, **f)**, **h)**, **j)**, **l)** and **n)**) of the BE-metaD simulations of chignolin. The title of each subplot indicates how the RCs of the BE-metaD simulations were obtained.

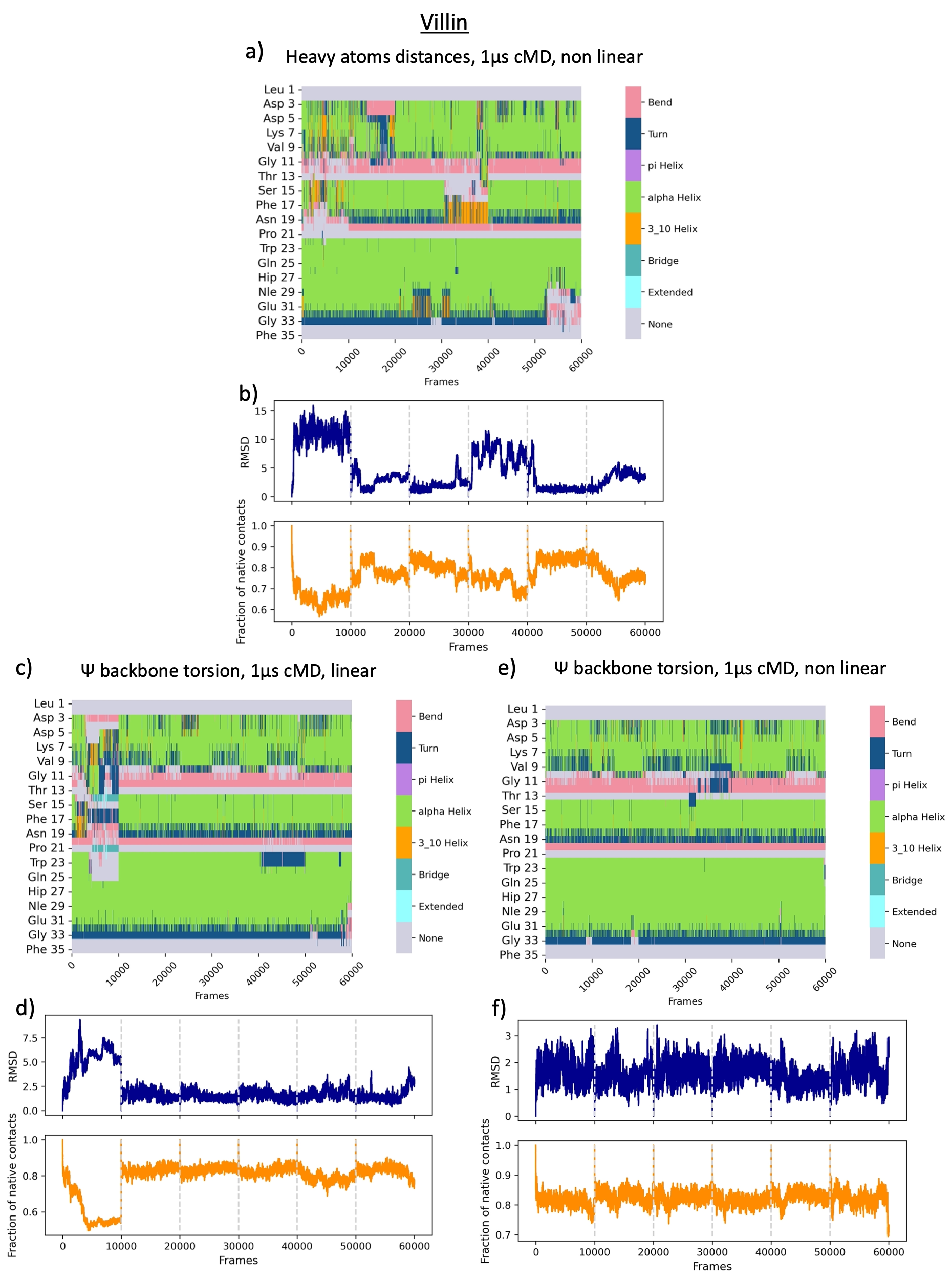

**Figure S3. Analyses of villin simulations.** Secondary structure (**a)**, **c)**, and **e)**), RMSD and native contacts analyses (**b)**, **d)** and **f)**) of the BE-metaD simulations of villin. The title of each subplot indicates how the RCs of the BE-metaD simulations were obtained.

|  | Angles | Distances |
| --- | --- | --- |
| Chignolin | 18 (sin and cos of 9 dihedrals) | 45 |
| Villin | 68 (sin and cos of 34 dihedrals) | 561 (20) |

**Table S1.** Number of features on which SPIB was trained.

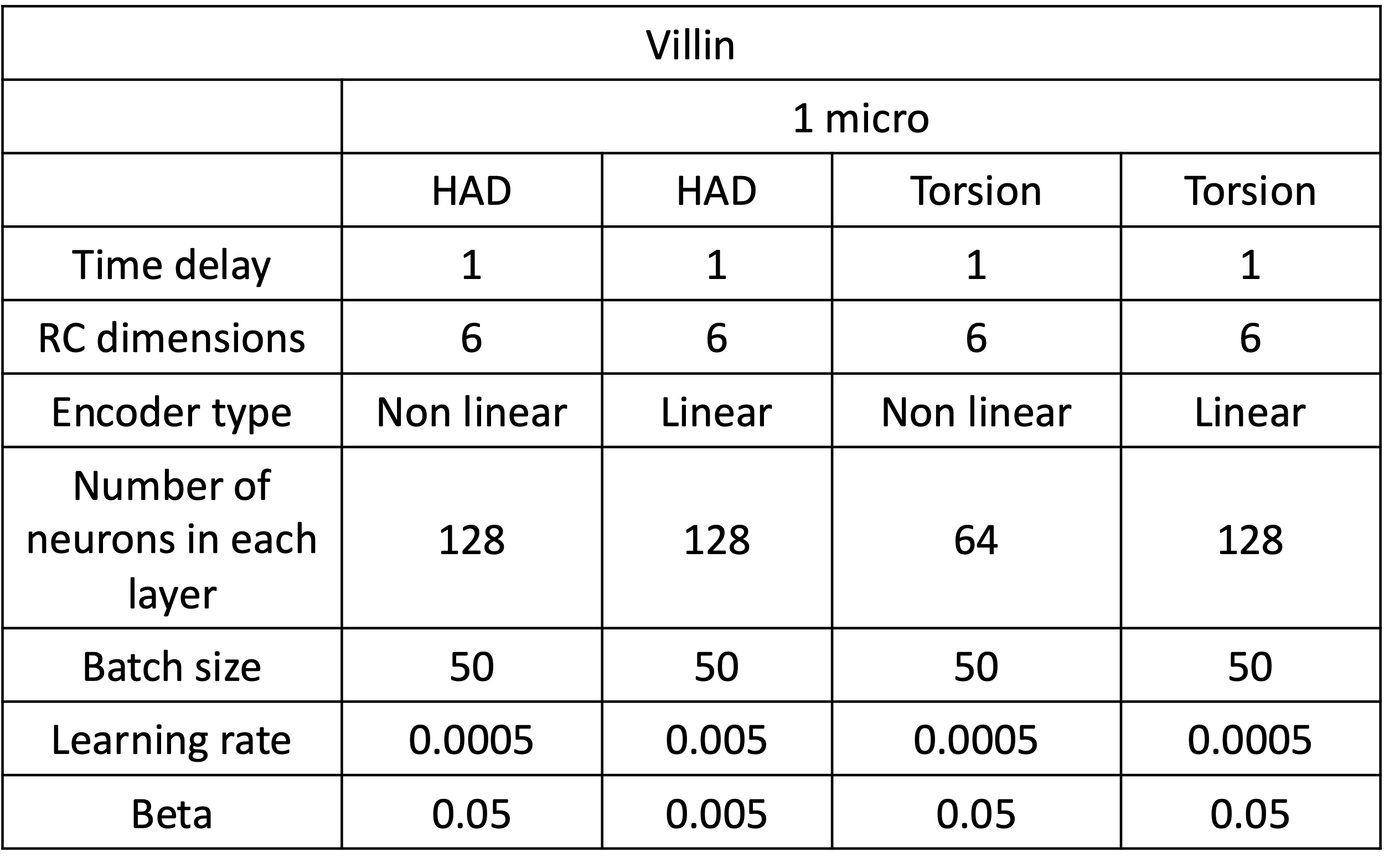

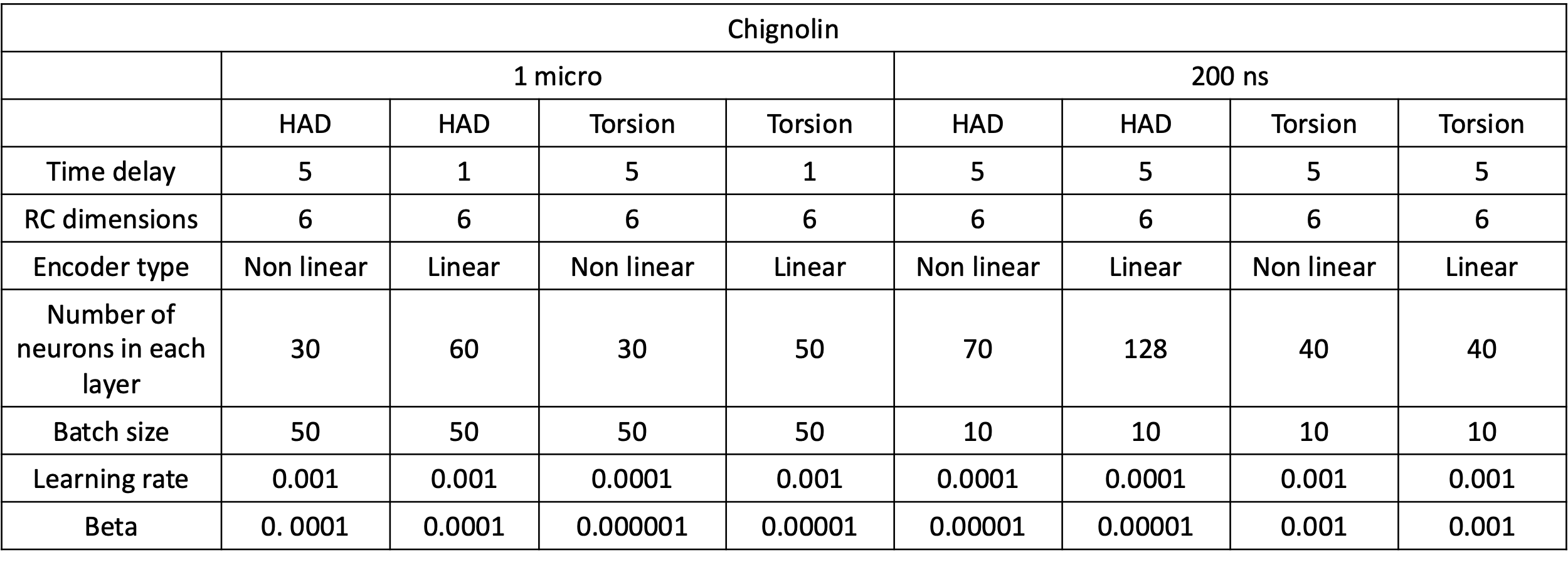
 **Table S2.** Hyperparameters for SPIB trainings
